# The enzymatic activity of the nsp14 exoribonuclease is critical for replication of Middle East respiratory syndrome-coronavirus

**DOI:** 10.1101/2020.06.19.162529

**Authors:** Natacha S. Ogando, Jessika C. Zevenhoven-Dobbe, Clara C. Posthuma, Eric J. Snijder

## Abstract

Coronaviruses (CoVs) stand out for their large RNA genome and complex RNA-synthesizing machinery comprising 16 nonstructural proteins (nsps). The bifunctional nsp14 contains an N-terminal 3’-to-5’ exoribonuclease (ExoN) and a C-terminal N7-methyltransferase (N7-MTase) domain. While the latter presumably operates during viral mRNA capping, ExoN is thought to mediate proofreading during genome replication. In line with such a role, ExoN-knockout mutants of mouse hepatitis virus (MHV) and severe acute respiratory syndrome coronavirus (SARS-CoV) were previously found to have a crippled but viable hypermutation phenotype. Remarkably, using an identical reverse genetics approach, an extensive mutagenesis study revealed the corresponding ExoN-knockout mutants of another betacoronavirus, Middle East respiratory syndrome coronavirus (MERS-CoV), to be non-viable. This is in agreement with observations previously made for alpha- and gammacoronaviruses. Only a single MERS-CoV ExoN active site mutant could be recovered, likely because the introduced D191E substitution is highly conservative in nature. For 11 other MERS-CoV ExoN active site mutants, not a trace of RNA synthesis could be detected, unless – in some cases – reversion had first occurred. Subsequently, we expressed and purified recombinant MERS-CoV nsp14 and established *in vitro* assays for both its ExoN and N7-MTase activities. All ExoN knockout mutations that were lethal when tested via reverse genetics were found to severely decrease ExoN activity, while not affecting N7-MTase activity. Our study thus reveals an additional function for MERS-CoV nsp14 ExoN, which apparently is critical for primary viral RNA synthesis, thus differentiating it from the proofreading activity thought to boost long-term replication fidelity in MHV and SARS-CoV.

**Importance:** The bifunctional nsp14 subunit of the coronavirus replicase contains 3’-to-5’ exoribonuclease (ExoN) and N7-methyltransferase (N7-MTase) domains. For the betacoronaviruses MHV and SARS-CoV, the ExoN domain was reported to promote the fidelity of genome replication, presumably by mediating some form of proofreading. For these viruses, ExoN knockout mutants are alive while displaying an increased mutation frequency. Strikingly, we now established that the equivalent knockout mutants of MERS-CoV ExoN are non-viable and completely deficient in RNA synthesis, thus revealing an additional and more critical function of ExoN in coronavirus replication. Both enzymatic activities of (recombinant) MERS-CoV nsp14 were evaluated using newly developed *in vitro* assays that can be used to characterize these key replicative enzymes in more detail and explore their potential as target for antiviral drug development.

## Introduction

RNA viruses commonly exhibit high mutation rates, a feature attributed to the relatively poor fidelity of their RNA-dependent RNA polymerase (RdRp) and the fact that nucleotide incorporation errors go uncorrected. This lack of proofreading contributes to the generation of ‘quasispecies’ populations, clouds of genome sequence variants that are subject to continuous natural selection (1–3). On the one hand, their genetic heterogeneity allows RNA viruses to rapidly adapt to changing circumstances, in order to overcome environmental challenges such as host switching, antiviral drug treatment, or host immune responses (4, 5). On the other hand, the accumulation of an excessive number of deleterious mutations can result in ‘error catastrophe’ and, consequently, in the extinction of a viral species (6–8). In order to balance these opposing principles, RNA viruses are thought to operate close to their so-called ‘error threshold’, while balancing the interdependent parameters of replication fidelity, genome size, and genome complexity (9, 10). This interplay is thought to have restricted the expansion of RNA virus genome size, which is below 15 kilobases (kb) for most RNA virus families (10–12).

The largest RNA virus genomes currently known are found in the order *Nidovirales*, which includes the coronavirus (CoV) family and also the recently discovered planarian secretory cell nidovirus (PSCNV; (12, 13)), which has the largest RNA genome identified thus far (41.1 kb). One of the molecular mechanisms potentially driving the unprecedented expansion of nidovirus genomes emerged about 17 years ago, during the in-depth bioinformatics analysis of the genome and proteome of the severe acute respiratory syndrome coronavirus (SARS-CoV). During this analysis, Alexander Gorbalenya and colleagues identified a putative 3’-to-5’ exoribonuclease (ExoN) signature sequence in the N-terminal domain of nonstructural protein 14 (nsp14), a subunit of the large replicase polyprotein encoded by CoVs and related large-genome nidoviruses. Strikingly, this ExoN domain was found to be lacking in the replicases of nidoviruses with small(er) genomes (specifically, arteriviruses), and therefore it was proposed that the enzyme may provide a form of ‘proofreading activity’ that could have promoted the expansion of large nidoviral genomes to their current size (10–12, 14). Comparative sequence analysis with cellular homologs classified the nidoviral/CoV ExoN domain as a member of the superfamily of DEDDh exonucleases, which also includes the proofreading domains of many DNA polymerases as well as other eukaryotic and prokaryotic exonucleases (15). These enzymes catalyze the excision of nucleoside monophosphates from nucleic acids in the 3’-to-5’ direction, using a mechanism that depends on two divalent metal ions and a reactive water molecule (16–18). Five conserved active site residues arranged in three canonical motifs (I, II and III; Fig. 1) orchestrate ExoN activity (14, 19–21). Additionally, the domain incorporates two zinc finger (ZF) motifs (10), ZF1 and ZF2 (Fig. 1), that were hypothesized to contribute to the structural stability and catalytic activity of ExoN, respectively (20).

**Figure 1.**
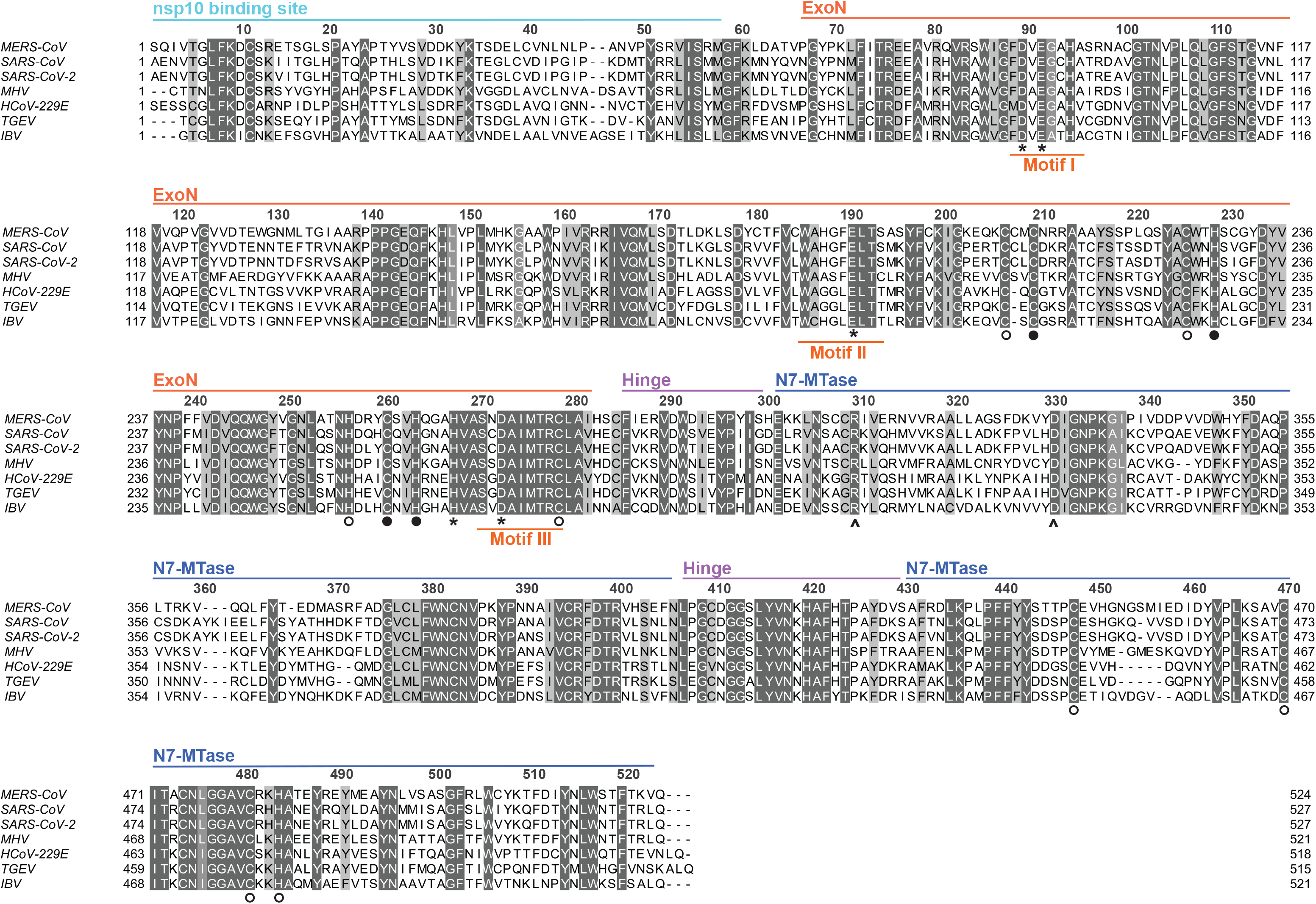
Alignment of nsp14 amino acid sequences from selected coronaviruses. Sequences of the ExoN and N7Mtase domains in MERS-CoV (NC-019843); SARS-CoV (NC_004718); SARS-CoV-2 (NC_045512.2); MHV (NP_045298); HCoV-229E (NC_002645); TGEV (AJ271965); and IBV (NP_040829) were used for the analysis. The different domains indicated on the top are based on the SARS-CoV-nsp14 secondary structure (PDB 5NFY (21)). Fully conserved residues are boxed in dark grey with white letters (above 70% conservation), whereas partially conserved residues are displayed in lighter shades of grey. Catalytic residues and residues involved in formation of zinc fingers are marked with asterisks and circles, respectively. Full circles indicate zinc fingers targeted by mutagenesis (Fig. 2A) while two black arrows identify the two N7Mtase domain residues mutated to generate the methyltransferase negative control used in biochemical assays. The alignment was generated using Clustal Omega (93) and edited using Jalview version 2.11 (94).

The predicted 3’-to-5’ exoribonuclease activity of the CoV ExoN domain was first confirmed *in vitro*, in biochemical assays using recombinant SARS-CoV nsp14 and different synthetic RNA substrates (19). Originally, residues D90/E92 (motif I), D243 (motif II), and D273 (motif III) were identified as putative active site residues of SARS-CoV nsp14 (14, 19). However, the SARS-CoV nsp14 crystal structure revealed E191 rather than D243 to be the acidic active residue in Motif II, demonstrating that ExoN is in fact a DEEDh enzyme (20). By using reverse genetics of the alphacoronavirus human coronavirus 229E (HCoV-229E), Minskaia *et al.* demonstrated that inactivation of the ExoN active site results in failure to recover infectious viral progeny (19).

Interestingly, a quite different phenotype was described for the corresponding ExoN-knockout mutants of two betacoronaviruses, mouse hepatitis virus (MHV) and SARS-CoV. While ExoN inactivation decreased replication fidelity in these viruses, conferring a ‘mutator phenotype’, the mutants were viable, both in cell culture (22, 23) and in animal models (24). These findings suggested that ExoN may indeed be part of an error correction mechanism. Subsequently, the ability of ExoN to excise 3’-terminal mismatched nucleotides from a double-stranded (ds) RNA substrate was demonstrated *in vitro* using recombinant SARS-CoV nsp14 (25). Furthermore, this activity was shown to be strongly enhanced (up to 35-fold) by the presence of nsp10, a small upstream subunit of the CoV replicase (26). The two subunits were proposed to operate, together with the nsp12-RdRp, in repairing mismatches that may be introduced during CoV RNA synthesis (21, 27). In cell culture, MHV and SARS-CoV mutants lacking ExoN activity exhibit increased sensitivity to mutagenic agents like 5-fluoracil (5-FU), compounds to which the wild-type virus is relatively resistant (28, 29). Recently, ExoN activity was also implicated in CoV RNA recombination, as an MHV ExoN knockout mutant exhibited altered recombination patterns, possibly reflecting its involvement in other activities than error correction during CoV replication and subgenomic mRNA synthesis (30). Outside the order *Nidovirales*, arenaviruses are the only other RNA viruses known to employ an ExoN domain, which is part of the arenavirus nucleoprotein and has been implicated in fidelity control (31) and/or immune evasion, the latter by degrading viral dsRNA (32, 33). Based on results obtained with TGEV and MHV ExoN knockout mutants, also the CoV ExoN activity was suggested to counteract innate responses (34, 35).

In the meantime, CoV nsp14 was proven to be a bifunctional protein by the discovery of an (N7-guanine)-methyltransferase (N7-MTase) activity in its C-terminal domain (36)(Fig. 1). This enzymatic activity was further corroborated *in vitro*, using biochemical assays with purified recombinant SARS-CoV nsp14. The enzyme was found capable of methylating cap analogues or GTP substrates, in the presence of S-adenosyl methionine (SAM) as methyl donor (36, 37). The N7-MTase was postulated to be a key factor for equipping CoV mRNAs with a functional 5’-terminal cap structure, as N7-methylation is essential for cap recognition by the cellular translation machinery (25). Although, the characterization of the nsp14 N7-MTase active site and reaction mechanism was not completed, alanine scanning mutagenesis and *in vitro* assays with nsp14 highlighted several key residues (Fig. 1)(36, 38, 39). Moreover, crystal structures of SARS-CoV nsp14 in complex with its nsp10 co-factor (PDB entries 5C8U and 5NFY) revealed several unique structural and functional features (Ma et al., 2015; Ferron et al., 2018). These combined structural and biochemical studies confirmed that the two enzymatic domains of nsp14 are functionally distinct (36) and physically independent (20, 21). Still, the two activities are structurally intertwined, as it seems that the N7-MTase activity depends on the integrity of the N-terminal ExoN domain, whereas the flexibility of the protein is modulated by a hinge region connecting the two domains (21).

Coronaviruses are abundantly present in mammalian reservoir species, including bats, and pose a continuous zoonotic threat (40–43). To date, seven CoVs that can infect humans have been identified, and among these the severe acute respiratory syndrome coronavirus 2 (SARS-CoV-2) is currently causing an unprecedented pandemic outbreak. The previous CoV to emerge, in 2012, was the Middle East respiratory syndrome coronavirus (MERS-CoV) (44). Due to zoonotic transfer from dromedary camels and subsequent nosocomial transmission, MERS-CoV still continues to circulate and cause serious human disease, primarily in the Arabian Peninsula (45). Occasional spread to other countries has also occurred, including an outbreak with 186 confirmed cases in South Korea in 2015 (46–48). Like SARS-CoV/SARS-CoV2 and MHV, MERS-CoV is classified as a member of the *betacoronavirus* genus, although it belongs to a different lineage (subgenus) of that cluster (49, 50). The current lack of approved therapeutics and vaccines to prevent or treat CoV infections, as well as the general threat posed by emerging CoVs, necessitates the further in-depth characterization of CoV replication and replicative enzymes. In this context, the quite different phenotypes described for ExoN knockout mutants of other CoVs (see above) prompted us to study the importance of this enzyme for MERS-CoV replication. To this end, using both reverse genetics and biochemical assays with recombinant nsp14, we engaged in an extensive site-directed mutagenesis study, targeting all active site residues of the MERS-CoV ExoN domain. Strikingly, in contrast to what was observed for other betacoronaviruses, our studies revealed that ExoN inactivation renders MERS-CoV RNA synthesis undetectable and results in failure to recover viable virus progeny. Our biochemical evaluation of nsp14 mutants suggests that this is not caused by inadvertent side-effects of ExoN inactivation on N7-MTase activity. Our combined data suggest that MERS-CoV ExoN and/or nsp14 play a more direct and fundamental role in CoV RNA synthesis than merely safeguarding the long-term fidelity of replication, and can be considered a prominent target for the development of antiviral drugs.

## Results

### ExoN inactivation is lethal for MERS-CoV

Previous studies into CoV ExoN function involved its biochemical characterization (based almost exclusively on the SARS-CoV version of the enzyme) and the phenotypic analysis of (predicted) ExoN knockout virus mutants, generated using reverse genetics approaches. The latter studies yielded replication-incompetent ExoN knockout mutants for the alphacoronaviruses HCoV-229E (19) and transmissible gastroenteritis virus (TGEV) (34). However, the equivalent mutants of the betacoronaviruses SARS-CoV and MHV-A59 were somewhat crippled but clearly viable, while displaying a 15-to 20-fold increased mutation rate (Eckerle, Lu et al. 2007, Eckerle, Becker et al. 2010). An alignment of CoV nsp14 amino acid sequences is presented in Fig. 1, including SARS-CoV-2, which emerged during the course of this project. It highlights the key motifs/residues of the two enzymatic domains of nsp14, as well as other structural elements, like the nsp10 binding site, the hinge region connecting the ExoN and N7-MTase domains, and three previously identified zinc finger domains (20, 21). The alignment also illustrates the generally high degree of nsp14 sequence conservation across different CoV (sub)genera.

In the present study, we targeted all five predicted active site residues of the MERS-CoV ExoN domain (D90, E92, E191, D273, and H268) by replacing them with alanine as well as more conservative substitutions (D to E or Q; E to D or Q). This yielded a total of 14 ExoN active site mutants (Fig. 2A), including the D90A/E92A motif-I double mutant (DM) that was frequently used in studies of the viable ExoN knockout mutants for MHV and SARS-CoV. A BAC-based MERS-CoV reverse genetics system (51, 52), based on the sequence of the EMC/2012 isolate of MERS-CoV (53), served as the starting point to evaluate our ExoN mutants by transfection of full-length RNA that was obtained by *in vitro* transcription using T7 RNA polymerase.

**Figure 2.**
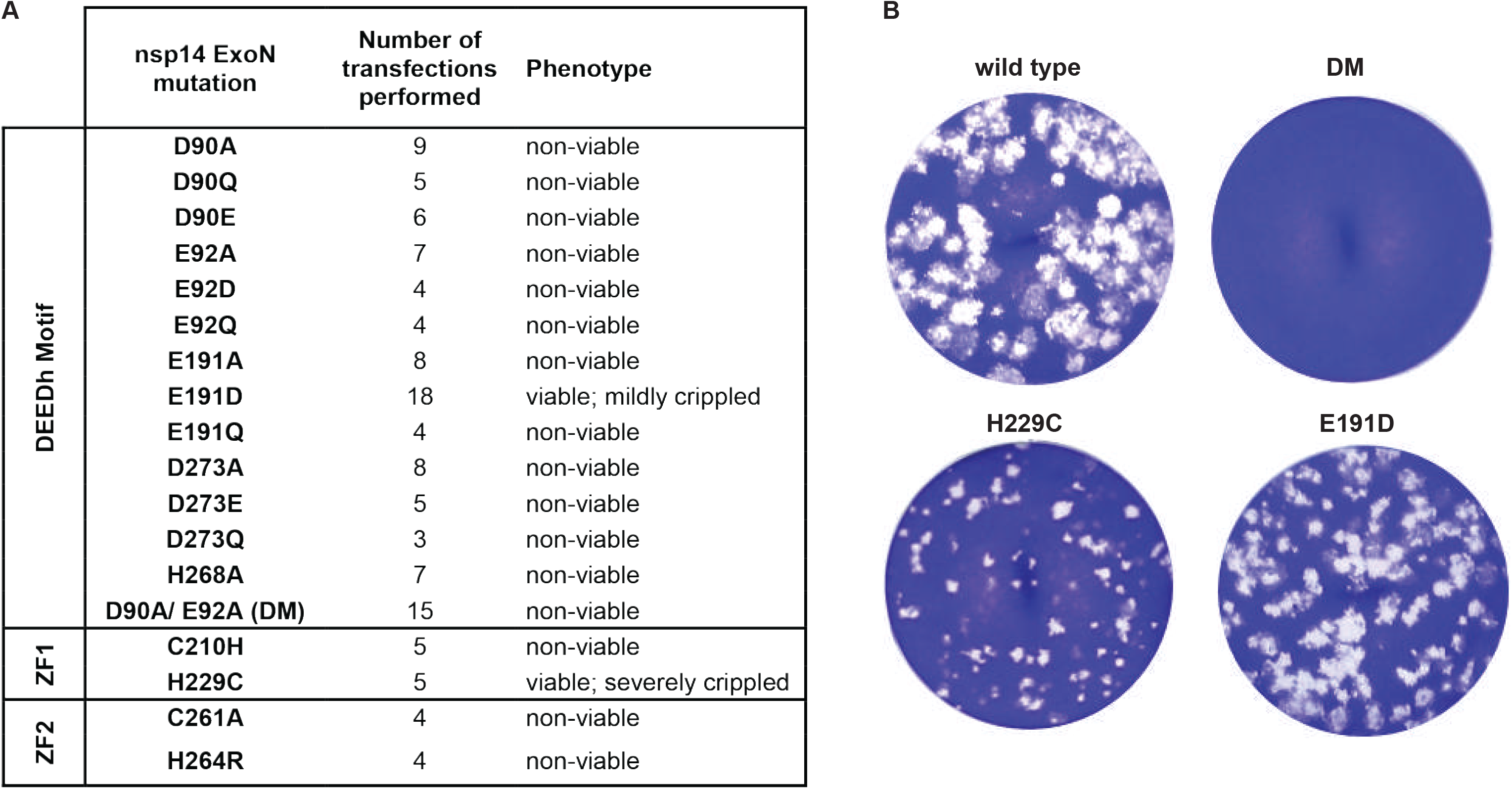
(A) Phenotype of MERS-CoV nsp14 ExoN mutants used in this study, scored at 2 d p.t.. (B) Comparison of plaque phenotype of selected ExoN mutants in HuH7 cells. Plaque assays were performed using supernatants harvested form transfected cells at 2 d p.t., which were diluted 10^−4^ for wt and mutant E191D mutant and used in undiluted form for the H229C and DM mutants.

Transcripts were electroporated into BHK-21 cells, which lack the DPP4 receptor required for natural MERS-CoV infection (Chan, Chan et al. 2013, Raj, Mou et al. 2013) but are commonly used to launch engineered CoV mutants because of their excellent survival of the electroporation procedure (19, 22, 23, 34, 51, 54). As BHK-21 cells have a severely compromised innate immune response (55), they would seem an appropriate cell line to launch ExoN knockout mutants also in case the enzyme would be needed to counter innate immunity (Becares, Pascual-Iglesias et al. 2016, Case, Li et al. 2018). To amplify any progeny virus released, transfected BHK-21 cells were mixed with either innate immune-deficient (Vero) or - competent (HuH7) cells, which both are naturally susceptible to MERS-CoV infection.

In stark contrast to what was previously described for MHV and SARS-CoV, mutagenesis of ExoN active site residues was found to fully abrogate MERS-CoV replication. When transfected cell cultures were analyzed using immunofluorescence microscopy at 2 days post transfection (d p.t.), abundant signal was always observed for wild-type MERS-CoV, but no sign of virus replication was observed for 13 out of 14 mutants tested (Fig. 2), regardless whether Vero or HUH7 cells were used for propagation of recombinant virus. Furthermore, infectious progeny was not detected when transfected cell culture supernatants were analyzed in plaque assays (Fig. 2 and data not shown). The single exception was the mutant carrying the conservative E191D replacement in ExoN motif II (Fig. 1), which was alive but somewhat crippled, as will be discussed in more detail below. These results were consistent across a large number of independent repeats (>10 for several of the mutants), performed with RNA transcribed from independently engineered (and fully sequenced) duplicate full-length cDNA clones. The non-viable phenotype of MERS-CoV ExoN mutants in both cell types suggests that innate immune responses did not influence the outcome of these experiments.

### ExoN inactivation abrogates MERS-CoV RNA synthesis

For a selection of MERS-CoV ExoN knockout mutants, intracellular RNA was isolated from transfected cell cultures after 48 h and analyzed by in-gel hybridization and RT-PCR to more rigorously verify the lack of viral RNA synthesis (Fig. 3). In this analysis, a non-viable MERS-CoV mutant with an in-frame 100-aa deletion in the nsp12-RdRp domain was used as a negative control (NC) for viral RNA synthesis, in order to assess and correct for the detection of any residual full-length RNA transcript that might still be present at this timepoint after transfection. Upon direct in-gel hybridization analysis using a ^32^P-labeled probe recognizing the 3’ end of all viral mRNAs, the characteristic nested set of MERS-CoV transcripts could only be detected for the E191D mutant and the wt control virus (Fig. 3). Even after a 28-day exposure of the phosphor imager screen (data not shown), signal could not be detected for any of the other mutants.

**Figure 3.**
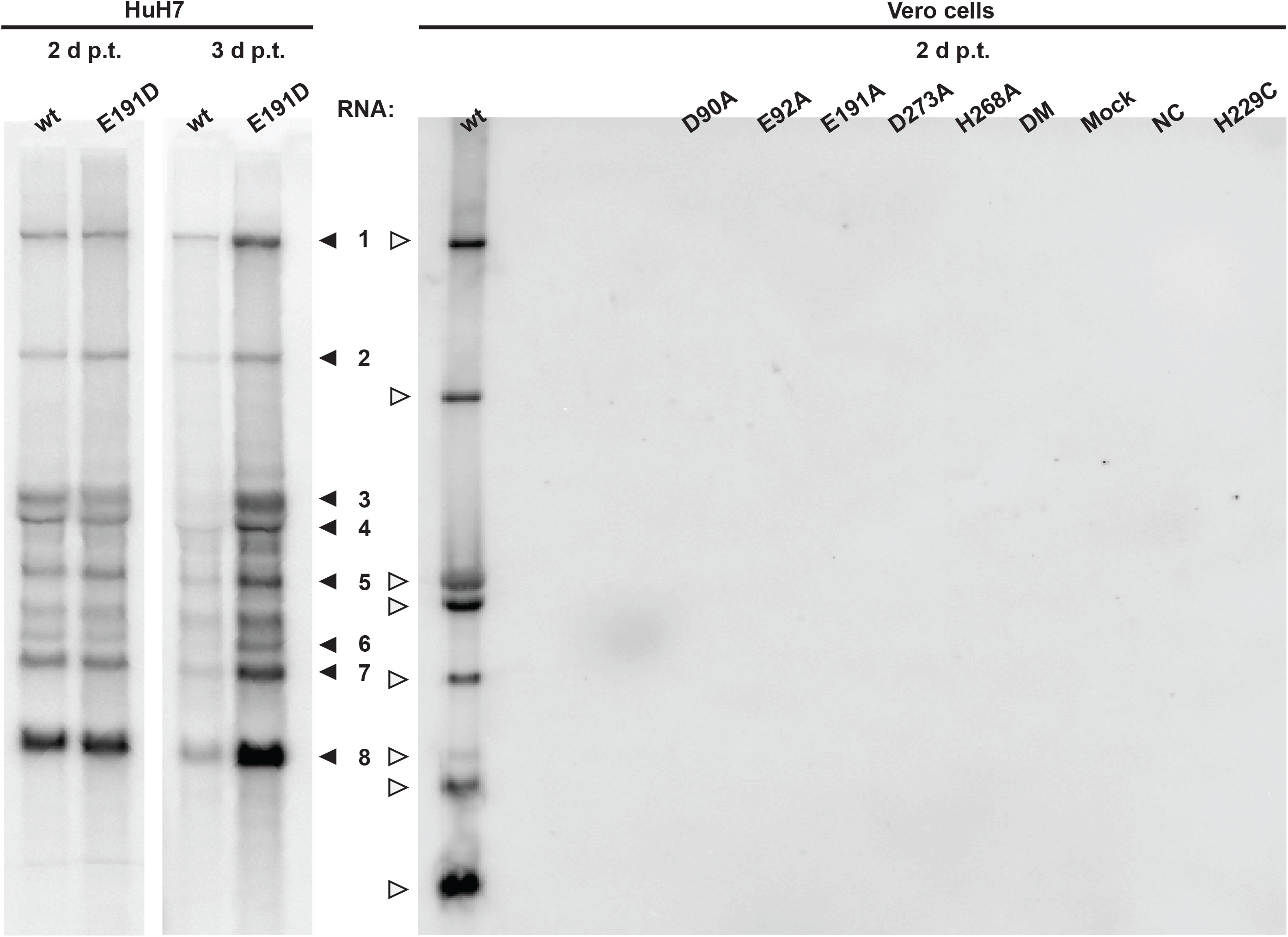
Impact of ExoN inactivation on intracellular RNA synthesis. In-gel hybridization analysis of intracellular RNA isolated after 2 or 3 days post transfection of transfected BHK-21 cells, which were subsequently mixed with HuH7 or Vero cells as indicated. Purified RNA was separated in an agarose gel and probed with a radiolabeled oligonucleotide probe recognizing the MERS-CoV genome and subgenomic mRNAs.

The lack of MERS-CoV-specific RNA synthesis was further analyzed using RT-PCR assays specifically detecting genomic RNA or subgenomic mRNA3. RNA accumulation was evaluated at 1 and 2 d p.t. for seven selected ExoN active site mutants (D90A, D90E, E191A, E191D, D273A, H268A, DM) using samples from two independent experiments both comprising duplicate transfections for each mutant. Again, MERS-CoV-specific genomic and subgenomic RNA synthesis was only detected for the E191D mutant and the wt virus control (data not shown). For all other mutants, the RT-PCR assays yielded Ct values equivalent to those obtained for samples from mock-infected cells and the replication-deficient NC mutant. In conclusion, with the exception of E191D (see below), all our engineered ExoN active site mutations abrogated viral RNA synthesis completely, suggesting that in the case of MERS-CoV the enzyme is indispensable for basic replication in cell culture.

### Characterization of rMERS-CoV-nsp14-E191D replication kinetics and 5-FU sensitivity

Among our ExoN active site mutants, only the E191D mutant yielded viable progeny (Fig. 2). The mutant appeared to be genetically stable as the substitution was preserved upon multiple consecutive passages in HuH7 or Vero cells (data not shown). Interestingly, the E191D mutation transforms the DEEDh catalytic motif into the DEDDh motif that is characteristic for members of the exonuclease family to which the CoV ExoN belongs (56). In fact, when comparing ExoN sequences from different nidovirus taxa (14, 19) the equivalent of E191 alternates between E and D (57), in line with the observation that this mutation is tolerated in MERS-CoV ExoN.

To characterize the E191D mutant in more detail, its replication and fitness in cell culture were analyzed. Full-length genome sequencing of passage 2 of the E191D mutant virus revealed that it had acquired two other mutations when compared with the recombinant wt control: a synonymous mutation in the nsp2-coding region (U→C at nt position 2,315) and a non-synonymous mutation (C→A at nt position 6,541) specifying an A1235D substitution in the betacoronavirus-specific marker (βSM) domain of nsp3, which has been predicted to be a non-enzymatic domain (58) and is absent in alpha- and delta-coronaviruses (59, 60). Thus, we assumed that any changes in viral replication were likely caused by the E191D mutation in nsp14 ExoN. The same virus stock was used to assess growth kinetics in HuH7 cells (Fig. 4B) and Vero cells (Fig. 4A), which were found to be very similar for wt and mutant virus. Still, the E191D mutant was found to be somewhat crippled, yielding smaller plaque sizes and somewhat lower progeny titers in HuH7 cells (Fig. 4 B-C), but not in Vero cells (Fig. 4A).

**Figure 4.**
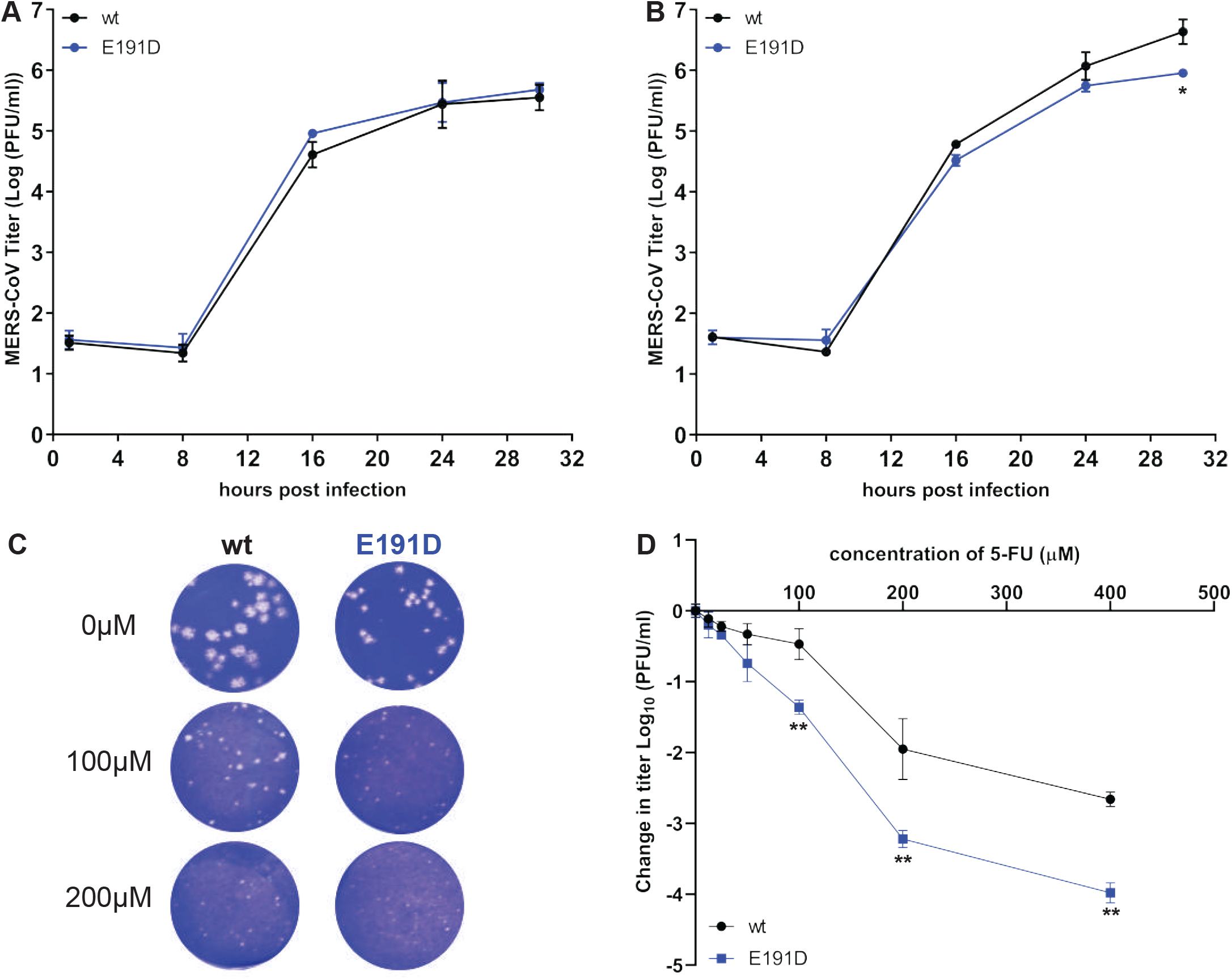
Characterization of growth kinetics of rMERS-nsp14-E191D and its sensitivity to 5-FU treatment. (A) Vero cells or (B) HuH7 cells were infected at an MOI of 3, supernatant was harvested at the indicated time points, and viral progeny titers were measured by plaque assay from two independent experiments using duplicates (n=4; mean ± sd is presented). (C) Plaque phenotype in HuH7 cells of rMERS-CoV nsp14-E191D and wt control in the absence or presence of the mutagenic agent 5-FU. (D) Dose response curve of wt and E191D mutant MERS-CoV in the presence of 5-FU concentrations up to 400μM (m.o.i. of 1; n=4; mean ± sd is presented). Statistical significance compared to wt at each time point (A and B) or concentration (D) was determined by paired t-test. All statistical analysis is denotated by asterisks: *, *P<0.05; ** p<0.005*.

We next examined the sensitivity of E191D and wt virus to the mutagenic agent 5-FU, which intracellularly is converted into a nucleoside analogue that is incorporated into viral RNA (61, 62). Previously, MHV and SARS-CoV ExoN knockout mutants were found to exhibit increased sensitivity to 5-FU treatment, in particular in multi-cycle experiments, which was attributed to a higher mutation frequency in the absence of ExoN-driven error correction (28). We employed this same assay to assess the phenotype of the E191D mutant in more detail, by performing plaque assays in HuH7 cells in the presence of increasing 5-FU concentrations (Fig. 4C) and by growing mutant and wt virus in the presence of increasing 5-FU concentrations (Fig. 4D). No cytotoxicity was observed in HuH7 cells following treatment with up to 400 μM 5-FU (data not shown).

Plaque assays in HuH7 cells were performed for 3 days, using a standard inoculum of 30 p.f.u. and an increasing amount of 5-FU in the overlay. Similar dose-dependent reductions of plaque size were observed for E191D and wt virus, with E191D plaques being barely visible upon treatment with 200 μM 5-FU (Fig. 4C). In an alternative experiment, Huh7 cells were infected with an m.o.i. of 1 and treated with an increasing 5-FU dose for 30 h, after which progeny virus titers were determined by regular plaque assay. Again, both viruses exhibited a similar concentration-dependent decrease of replication (Fig. 4D), although the E191D mutant appeared to be somewhat more sensitive to the mutagenic agent, yielding ~1-log lower progeny titers than wt virus upon treatment with 5-FU concentrations between 100 and 400 μM. These experiments demonstrated thatE191D and wt MERS-CoV are comparably sensitive to 5-FU treatment, suggesting that ExoN functionality is not strongly affected by the E191D substitution.

### ExoN zinc finger motifs are important for viral replication

Studies addressing the structural biology and biochemistry of SARS-CoV nsp14 indicated that the two ZF motifs within the ExoN domain contribute to either its structural stability (ZF1) or to catalytic activity (ZF2) (20, 21). Moreover, mutagenesis studies of the MHV and TGEV ZF1 domain supported their importance for viral replication in cell culture (34, 54). To study the impact of similar mutations on MERS-CoV replication, the nsp14 ZF1 and ZF2 domains were targeted with two mutations each and their impact on virus viability was evaluated as described above. Two ZF1 residues (C210 and H229) were mutated from H to C or vice versa, which could theoretically preserve the zinc-coordinating properties (63, 64). Two residues of the non-classical ZF2 motif were also substituted (C261A and H264R) to evaluate the same ZF mutations previously analyzed by Ma *et al.*, leading to disruption of ExoN activity *in vitro* (20).

The four ZF virus mutants were launched as described above, after which a low level of replication could be observed only for the H229C ZF1 mutant, for which the 2 d p.t. harvest yielded very small plaques and low progeny titers (Fig. 2B). For this mutant, RNA synthesis could not be detected by hybridization analysis (Fig. 3), but synthesis of genomic and subgenomic RNA (mRNA3) was detected by RT-PCR in intracellular RNA samples harvested at 2 d p.t. (data not shown). A 6 d p.t. harvest was used for full genome sequencing by NGS, which confirmed the presence of the engineered nsp14 mutation in addition to the appearance of some minor genetic variants (point mutations representing less than 15% of the total population) in different regions of the genome, including the ORF1a domain encoding for nsp3, nsp6, nsp8, and nsp9. Taken together, our observations defined the severely crippled phenotype of the H229C mutant. In combination with the fact that the other ZF mutations (C210H in ZF1, and C261A and H264R in ZF2) abolished all detectable MERS-CoV replication, our study establishes the importance of both ExoN ZF motifs for MERS-CoV viability.

### Development of a MERS-CoV ExoN activity assay using recombinant nsp14

In order to assess the impact of mutations on nsp14’s enzymatic activities, we set out to purify recombinant MERS-CoV nsp14 and develop an *in vitro* ExoN assay. Thus far, such an assay had only been described for the equivalent SARS-CoV protein (19, 20, 26, 65). Wild-type and mutant MERS-CoV nsp14 proteins carrying an N-terminal His-tag were expressed in *E. coli* Rosetta (DE3) pLysS. Proteins were purified by immobilized metal affinity chromatography (IMAC) followed by size exclusion chromatography. Upon SDS-PAGE, the purified MERS-CoV nsp14 was consistently detected as a doublet (with the lower band being most abundant), migrating at the expected molecular mass of ~55 kDa (Fig. 5). As a positive control, we purified SARS-CoV nsp14 (26) and used it during optimization of the enzymatic assays for MERS-CoV nsp14. The substrate used for ExoN activity assays was a 5’-^32^P-labeled 22-nucleotide (nt) long synthetic RNA, as previously used in similar assays with SARS-CoV nsp14 (referred to as oligonucleotide H4 in (26)). Nucleotides 5-22 of this substrate are predicted to form a hairpin with a stem consisting of seven G-C base pairs and a loop of 4 A’s (26), while the remaining 4 nucleotides form a 5’-terminal single-stranded tail.

**Figure 5.**
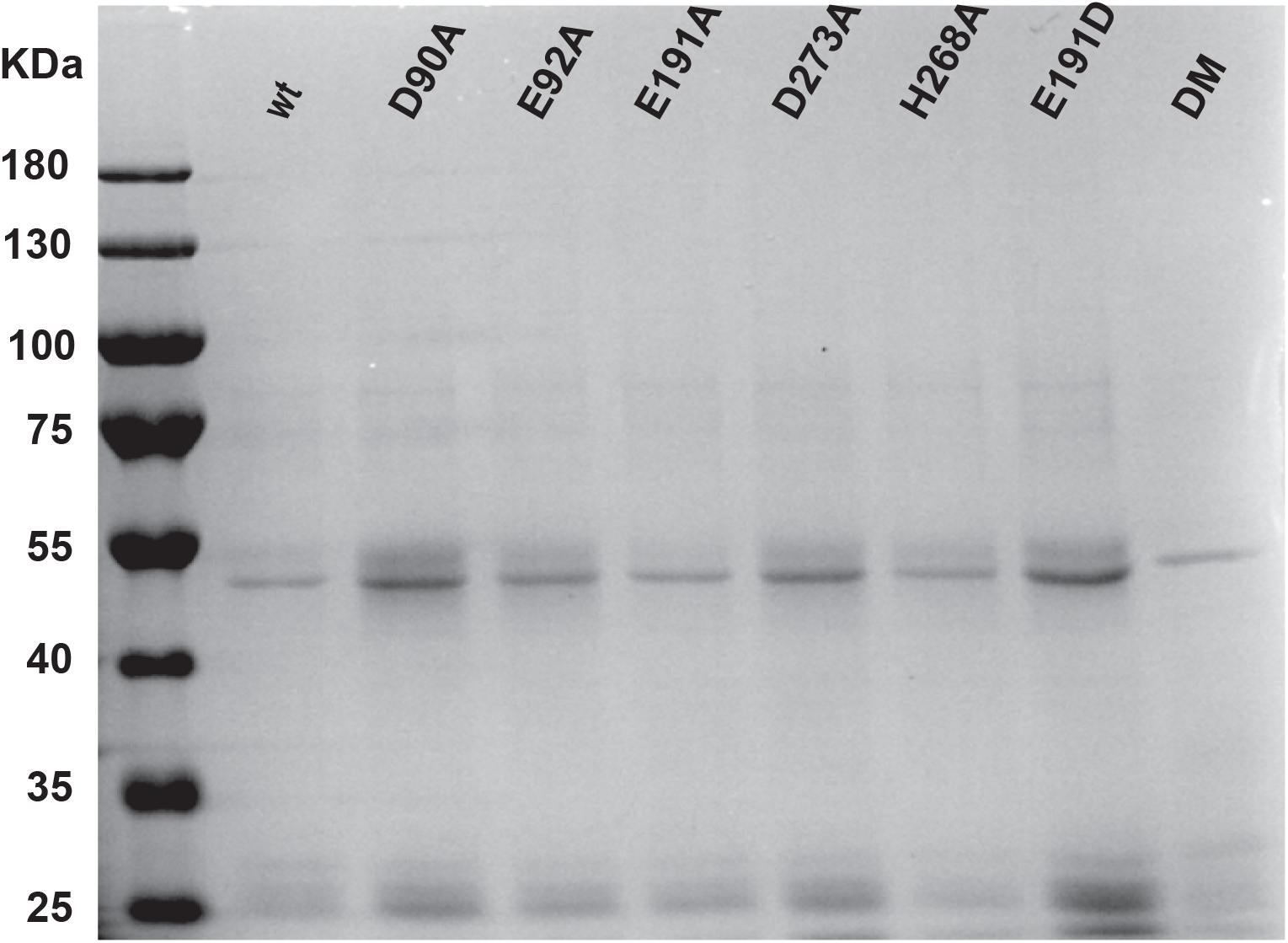
Expression and purification of recombinant MERS-CoV nsp14. N-terminally His-tagged wt and mutant MERS-CoV nsp14 (~55 kDa) was expressed in *E. coli*, affinity purified, and analyzed in a 10% SDS-PAGE gel that was stained with Coomassie Blue. The molecular masses of the protein marker (Invitrogen) are given in kDa.

Previously, the ExoN activity of SARS-CoV nsp14 was found to be dramatically stimulated by the addition of nsp10 as co-factor (26). Consequently, we also expressed and purified MERS-CoV nsp10 and optimized the ExoN assay by testing different molar ratios between nsp14 and nsp10 (Fig. 6A, left-hand side), different nsp14 concentrations (Fig. 6B, left-hand side), and by different incubation times (Fig. 7, left-hand side). MERS-CoV nsp14 ExoN activity was found to be stimulated by nsp10 in a dose-dependent manner (Fig. 6A), while nsp10 did not exhibit any nuclease activity by itself (Fig. 6B, nsp10 lane). The full-length substrate is more completely degraded when a fourfold (or higher) excess of nsp10 over nsp14 was used compared to the effect of merely increasing the nsp14 concentration in the assay (Fig. 6B). Similar observations were previously reported for SARS-CoV nsp14 (20, 26). Introduction of the D90A/E92A motif-I double substitution resulted in a major reduction of ExoN activity, although a certain level of residual activity was observed, in particular when using large amounts of nsp14 (Fig. 6B, right-hand side) or a relatively high nsp10:nsp14 ratio (Fig. 6A, right-hand side). Similar observations were previously made for SARS-CoV nsp14 (Bouvet, Imbert et al. 2012, Ma, Wu et al. 2015).

**Figure 6.**
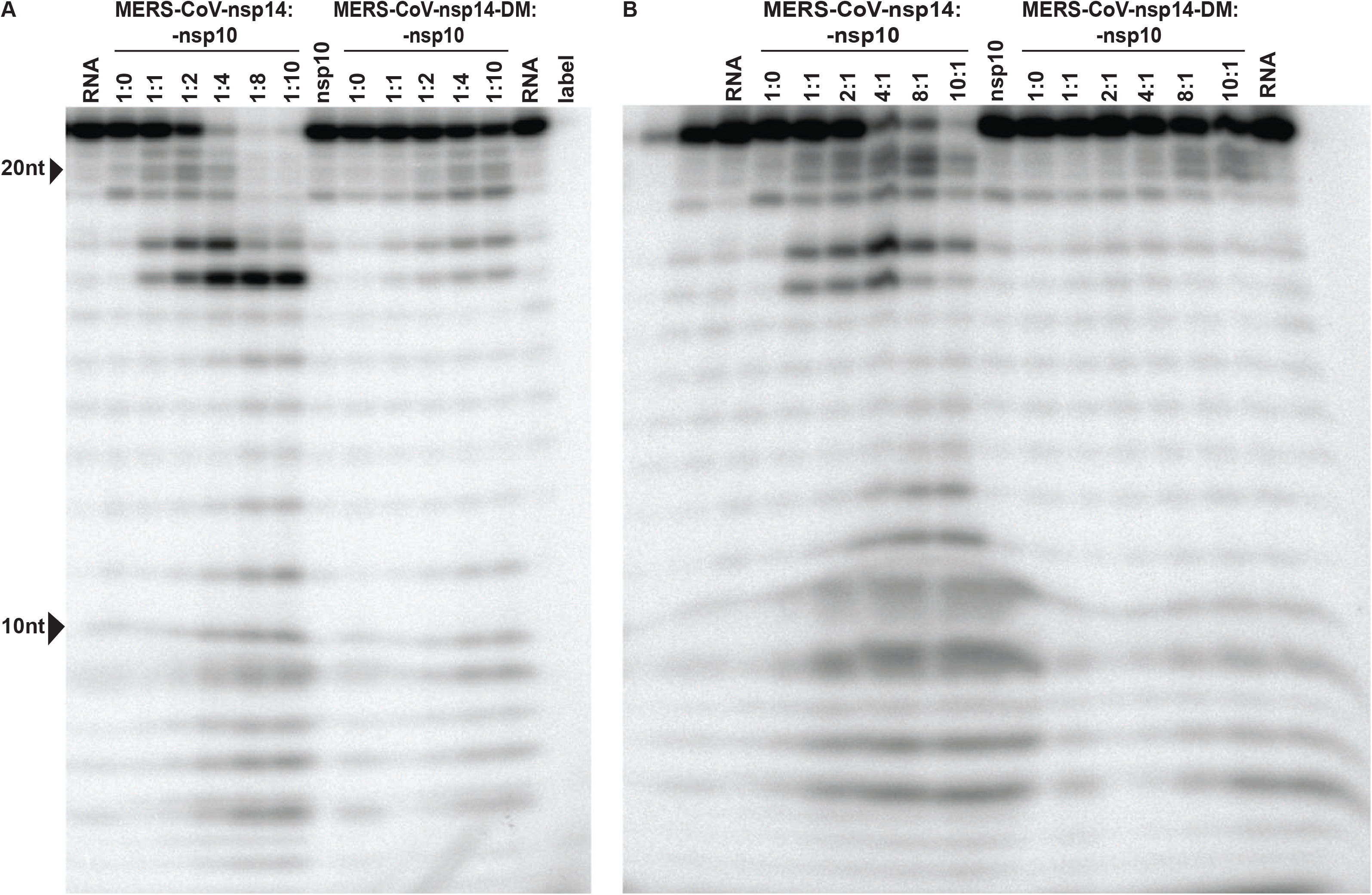
Optimization of MERS-CoV nsp14 *in vitro* ExoN assay conditions. The substrate for the assay was a 22-nt long synthetic RNA (H4) that was ^32^P-labeled at its 5’ terminus (*p-RNA). Cleavage products were separated by polyacrylamide gel electrophoresis and visualized by autoradiography. (A) Analysis of ExoN activity in the presence of an increasing amount of nsp10, using wt MERS-CoV-nsp14 (left) and the ExoN double knockout mutant (DM, D90A/E92A; right). The RNA substrate was hydrolyzed for 90 min at 37⁰C using a fixed concentration of nsp14 (200 nM) and an increasing nsp10 concentration, ranging from 0 to 1600 nM. (B) Evaluation of the ExoN activity of an increasing concentration (200 to 2000 nM) of wt or DM nsp14 in the presence of a fixed amount of nsp10 (200 nM).

**Figure 7.**
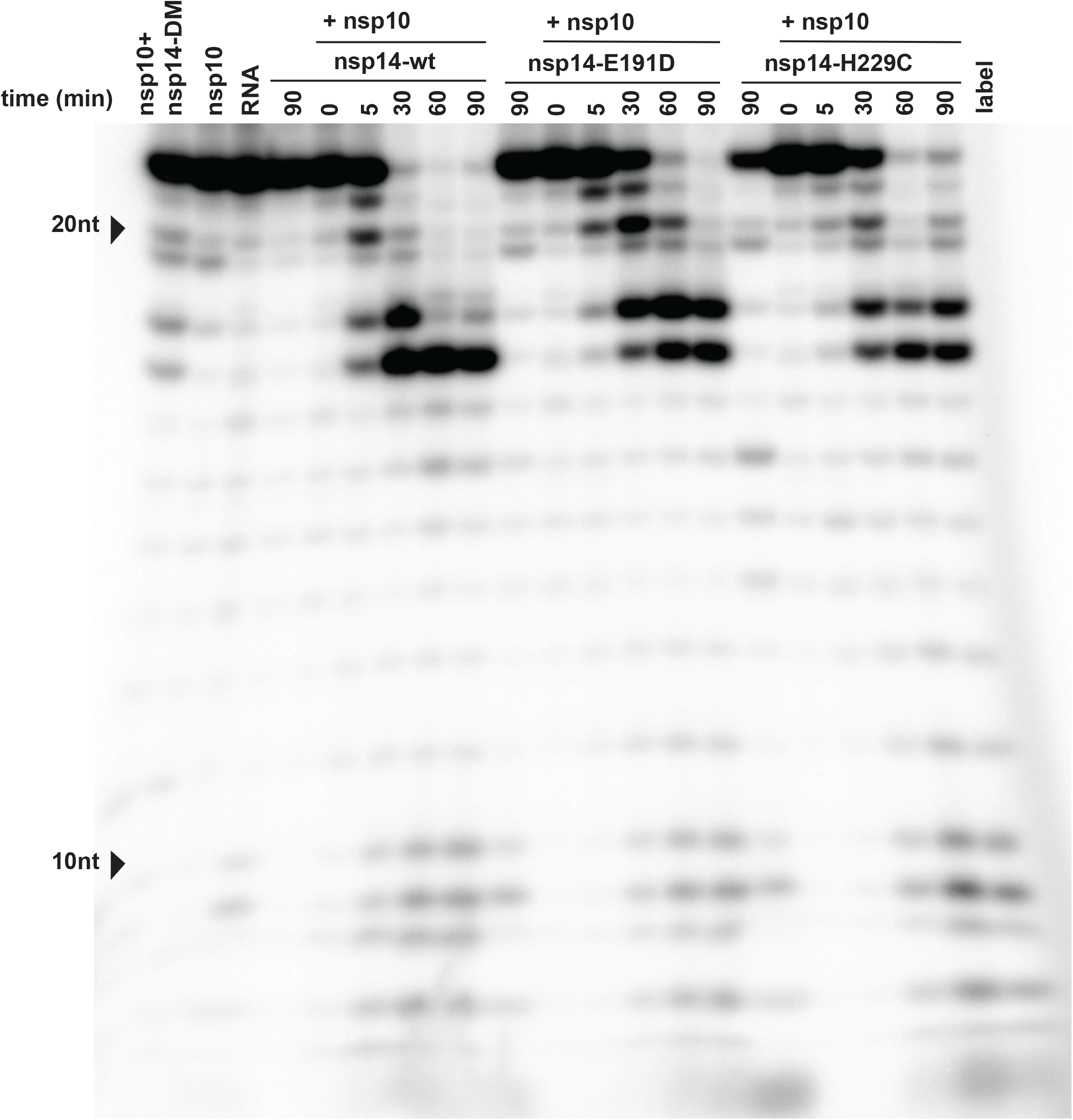
Time course analysis of the *in* vitro ExoN activity of MERS-CoV nsp14. The ExoN activity of different recombinant nsp14 proteins (wt, D90A/E92A, E191D and H229C) was evaluated by incubating 200 nM of nsp14 and 800 nM of nsp10 for 0, 5, 30, 60 and 90 min at 37⁰C. As controls, individual proteins (800 nM) were incubated for 90 min. For technical details, see the legend to Fig. 6.

Using a 4:1 ratio of nsp10 versus nsp14, MERS-CoV ExoN activity was analyzed in a time-course experiment, (Fig. 7). Over time, the full-length substrate was progressively converted to a set of degradation products in the size range of 6-18 nt. We anticipated that the structure of the H4 RNA substrate would change from a duplexed to a single-stranded conformation, upon digestion of one side of the hairpin’s stem by ExoN’s nuclease activity. As the ExoN enzymes of other CoVs were reported to prefer dsRNA substrates (19, 65), the degradation of the substrate might be slowed down substantially after removal of the first couple of nucleotides from its 3’ end (26). This would explain the abundant degradation products of 16 and 17 nt in length that were observed (Fig. 6-8), suggesting that the preference for dsRNA substrates is indeed shared by MERS-CoV ExoN.

Degradation of the substrate could be observed within 5 min and the full-length substrate was essentially gone after 30 min. A similar reaction with the nsp14 DM mutant resulted in only a small amount of substrate degradation after 90 min (Fig. 7, leftmost lane). Taken together, our results convincingly demonstrate the *in vitro* 3’-to-5’ exonuclease activity of purified MERS-CoV nsp14. As in the case of the SARS-CoV enzyme, nsp10 is a critical co-factor that can strikingly upregulate MERS-CoV ExoN activity *in vitro*.

### MERS-CoV nsp10 modulates nsp14 ExoN activity

In order to investigate differences that might explain the variable phenotype of CoV ExoN knockout virus mutants, we compared ExoN activities between SARS-CoV and MERS-CoV nsp14, using the optimized *in vitro* assay described above. An incubation time of 90 min was used in all experiments, unless indicated otherwise The nsp14 and nsp10 preparations of both viruses were first tested individually in an assay containing the H4 RNA substrate and Mg^+2^ ions (26, 66, 67). As expected, this revealed only traces of exonuclease activity for both nsp14 proteins (Fig. 8, lanes 4 and lane 6). When the two proteins were combined in the same reaction, a strong increase of ExoN activity was observed for both nsp14-nsp10 pairs, with the SARS-CoV pair appearing to be somewhat more processive than the MERS-CoV pair (Fig. 8, lanes 2 and 9).

**Figure 8.**
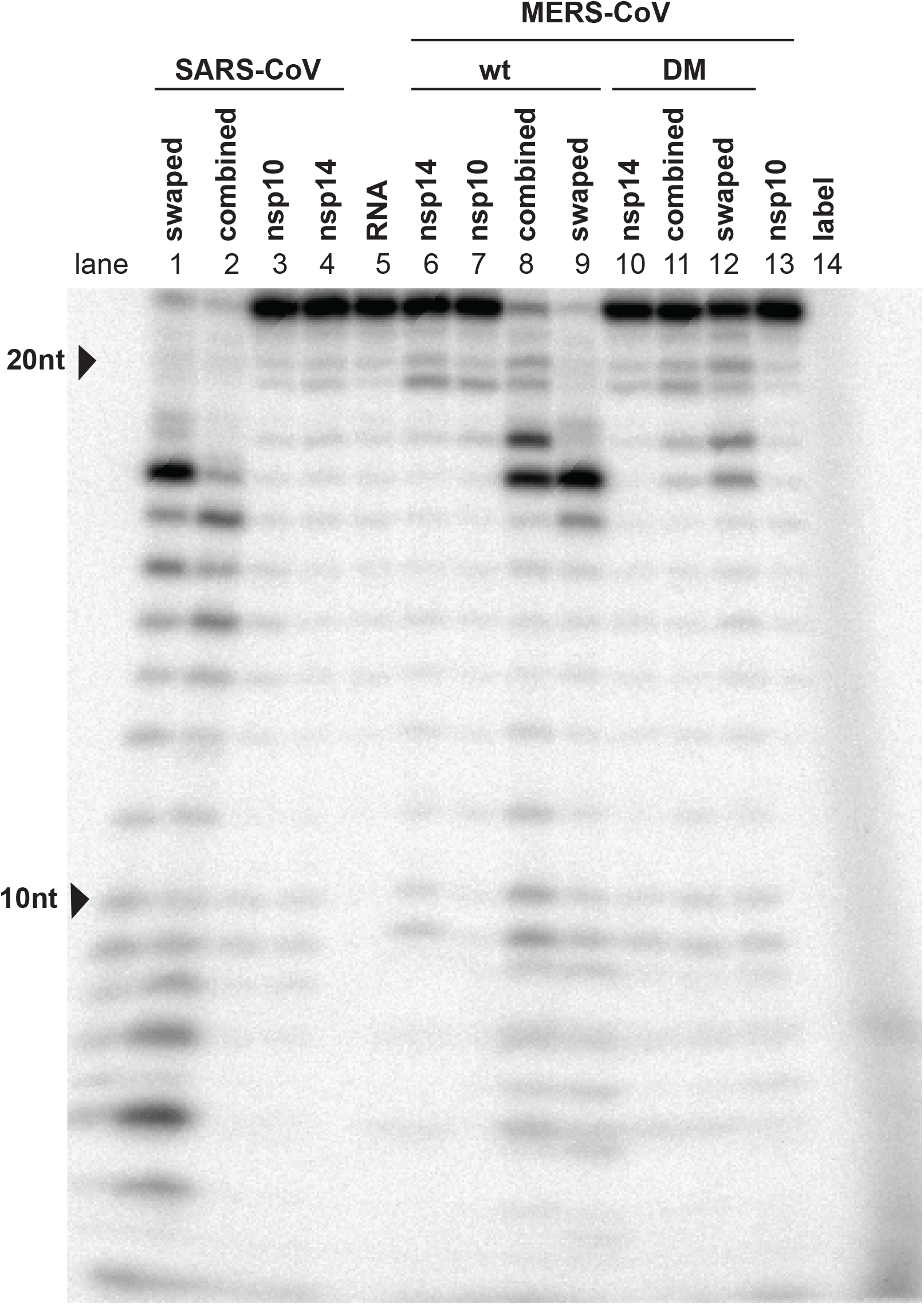
Cross-activation of the *in vitro* activity of SARS-CoV and MERS-CoV nsp14 by heterologous nsp10. The nsp10 co-factor was exchanged in ExoN assays performed with MERS-CoV and SARS-CoV nsp14, using a 1:4 ratio between nsp14 and nsp10 and a 90-min incubation at 37⁰C. For technical details, see the legend to Fig. 5.

The exchange of the SARS-CoV and MERS-CoV nsp10 co-factors revealed that they can cross-activate the ExoN activity of nsp14 from the other virus, although some changes in the pattern of degradation products were observed (Fig. 8, Lanes 1-2 and lanes 8-9). However, the residual ExoN activity of the motif I double mutant (DM) apparently was not affected by the choice of nsp10 co-factor (Fig. 8, compare lanes 11 and 12). The observed subtle changes in degradation product patterns are another indication that nsp10 modulates nsp14 ExoN activity, presumably using interaction surfaces that are well-conserved across CoV genera (66, 67).

### ExoN activity of MERS-CoV active site and H229C mutants

Having established the optimal conditions for MERS-CoV ExoN *in vitro* activity, we evaluated the impact of a subset of the DEEDh active site mutations that were used during our reverse genetic analysis (Fig. 2A). For each mutant tested, two protein batches were purified and analyzed independently in duplicate using the same batch of MERS-CoV nsp10 for all assays. As can be seen in Fig. 9, replacement with Ala of each of the five active site residues resulted in a near-complete loss of ExoN activity, with the D90A, E92A, and H268A substitutions appearing to be slightly less detrimental than E191A and motif III D273A. A clearly different result was again obtained with the E191D mutant, which displayed an activity level comparable to that of wt nsp14, corresponding with the properties of the corresponding virus mutant (Fig. 4). Overall, the severe impact of active site mutations on ExoN activity was fully in line with the non-viable phenotypes observed for the same mutants when tested using reverse genetics (Fig. 2).

**Figure 9.**
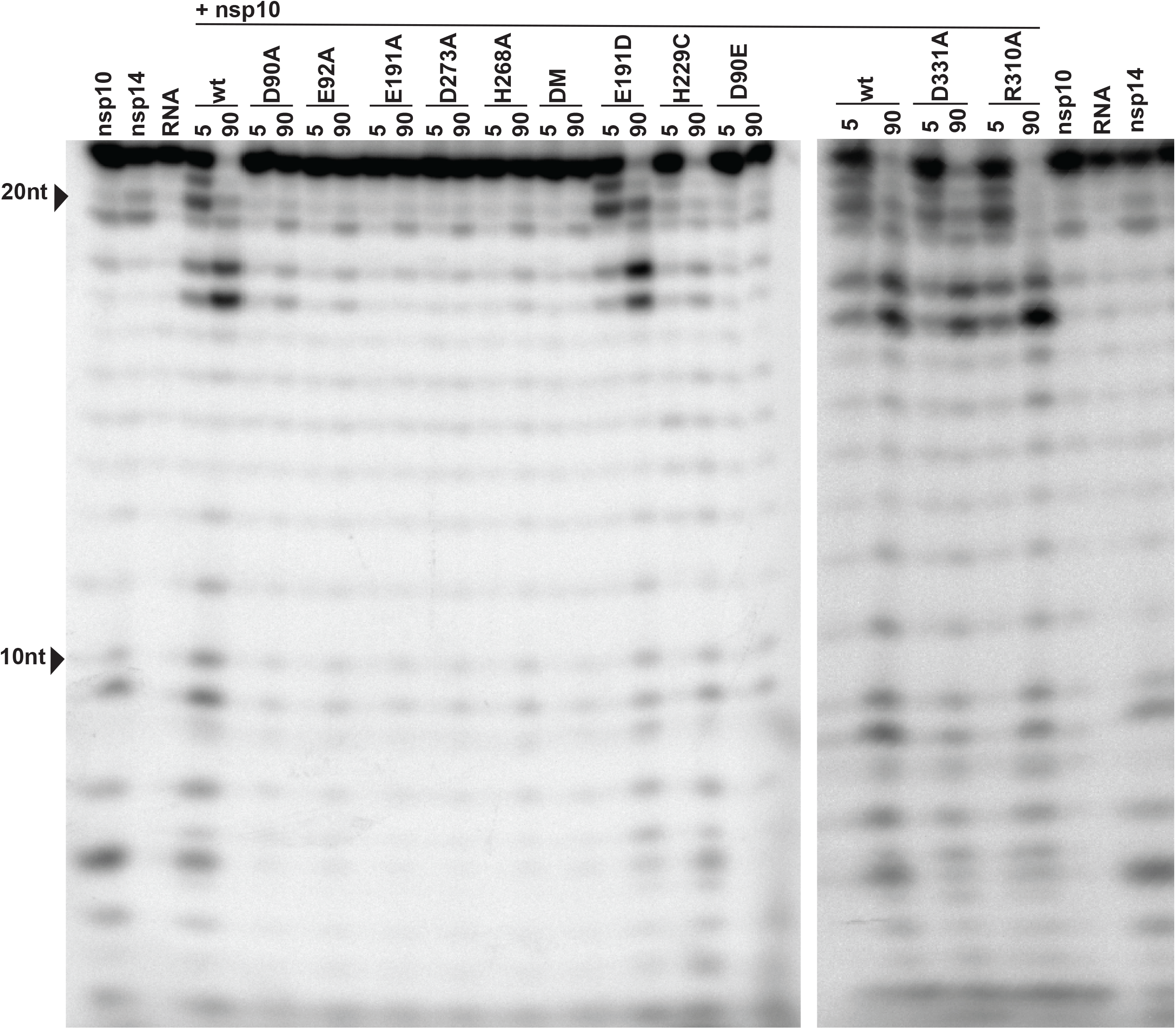
*In vitro* ExoN activity of MERS-CoV nsp14 mutants. Residues from the DEDDh catalytic motif and ZF1 motif of the nsp14 ExoN domain and the nsp14 N7MTase domain were mutated as indicated. Assays were performed using a 1:4 ratio between nsp14 and nsp10 and a 90-min incubation at 37⁰C. For technical details, see the legend to Fig. 6.

We also evaluated the impact of the H229C ZF1 mutation, which – despite its conservative nature - yielded a crippled mutant virus (Fig. 2) and of two N7-MTase mutations (discussed below). The N7-MTase mutants displayed wt nsp14-like ExoN activities (Fig. 9), suggesting that – as in SARS-CoV nsp14 - ExoN and N7-MTase activities are functionally separated (36). Analyzing the substrate degradation pattern of the H229C mutant (Fig. 7), the enzyme seems to be somewhat crippled when compared to wt nsp14. This suggests that this mutation alters ExoN activity *in vitro*, potentially by affecting the structure of the ExoN domain, as ZF1 is in close proximity of the nsp10 interaction surface (20). However, a similar reduction of ExoN activity was observed for the E191D mutant (Fig. 7), which was much more viable than the H229C mutant in the context of our reverse genetics studies. This suggests that the H229C replacement may affect additional functions or interactions of the ExoN domain that are important for viral RNA synthesis and viability.

### ExoN mutations do not interfere with N7-MTase activity *in vitro*

The nsp14 N7-MTtase activity is deemed essential for formation of a functional RNA cap, enabling the translation of CoV mRNAs and protecting them from degradation. Consequently, at least in theory, ExoN mutations could also be detrimental for virus replication if they would somehow affect the crucial enzymatic activity of the other nsp14 domain. In order to evaluate this possibility, the same recombinant protein preparations used in the ExoN assays (Fig. 9) were evaluated in an N7-MTase biochemical assay using the synthetic cap analogues GpppA and m7GpppA (control) as substrates. Moreover, nsp14 mutants R310A and D331A were used as negative controls in view of their predicted involvement in the binding of the triphosphate moiety of the RNA chain and the methyl donor (S-adenosylmethionine; SAM), respectively (20, 26, 36). In this assay, nsp14 can methylate GpppA by transferring the [3H]CH3 moiety provided by [^3^H]SAM. The resulting radio-labelled m7GpppA product can be quantified using a DEAE filter-binding assay, followed by liquid scintillation counting, and data normalization against the activity of wt control protein (25).

Recombinant MERS-CoV nsp14 was found to methylate GpppA, but not m7GpppA (Fig. 10A), which yielded a signal that was similar to the background signal in assays lacking nsp14 or substrate (data not shown). Methylation increased with time until reaching a plateau after 120 min (Fig. 10B). The N7-MTase activity of the various nsp14 mutants was compared with that of wt nsp14 after reaction times of 30 and 120 min (Fig. 10C). While the R310A and D331A control mutations fully inactivated the N7-MTase activity of MERS-CoV nsp14, none of the ExoN active site mutations tested was found to alter the enzyme’s activity. These results again support the notion that ExoN and N7-MTase domains are functionally separated, as previously demonstrated for SARS-CoV-nsp14 (36). We therefore conclude that the lethal impact of ExoN inactivation on MERS-CoV replication (Fig. 2A) cannot be attributed to inadvertent effects on the activity of the N7-MTase domain that is present in the same nsp14 replicase subunit.

**Figure 10.**
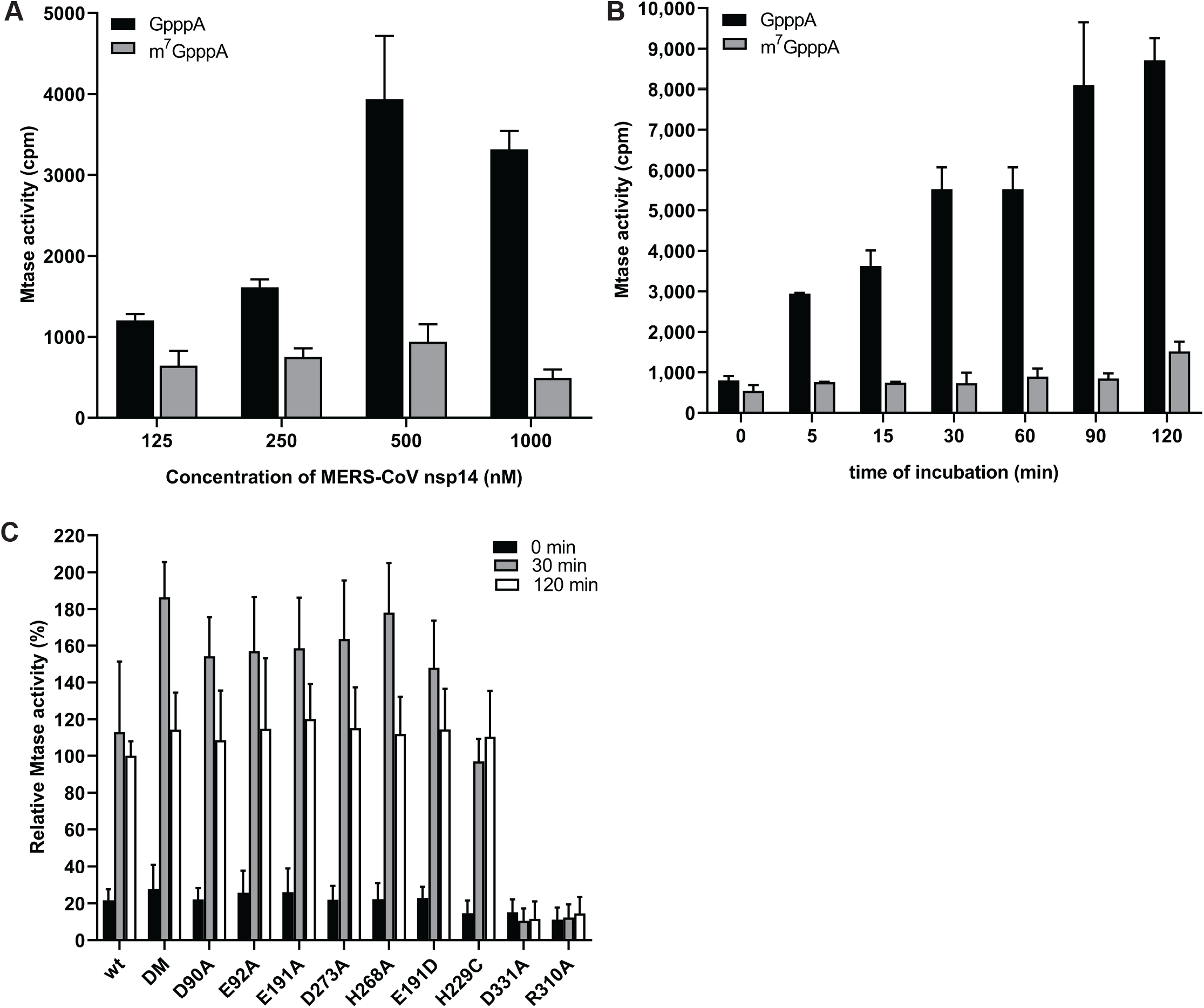
*In vitro* N7-MTase activity of MERS-CoV nsp14 mutants. The N7-MTase activity of recombinant nsp14 was analyzed *in vitro* by filter binding assay using synthetic cap analogues as substrate. (A) Increasing concentrations of MERS-CoV nsp14 were incubated with GpppA and m^7^GpppA in the presence of [^3^H]SAM for 30 min. (B) The ability of nsp14 to methylate GpppA or m^7^GpppA was determined after reaction times between 0 and 120 min at 30⁰C. (C) The ability of nsp14 mutants to methylate GpppA was measured in four times in duplicate. Values were normalized to the wt control (n=8; mean ± sd are shown).

## Discussion

In this study of MERS-CoV nsp14, we demonstrate that the impact of ExoN inactivation on virus viability and RNA synthesis distinguishes MERS-CoV from two other betacoronaviruses, MHV and SARS-CoV. Whereas ExoN inactivation in the latter two viruses yields viable mutants that are only mildly crippled and exhibit a “mutator phenotype” (22, 23, 29), both conservative and alanine substitutions of MERS-CoV ExoN catalytic residues abolished all detectable viral RNA synthesis (Fig. 3) and the release of viral progeny (Fig. 2). The only exception was the conservative E191D mutant, which was found to exhibit near-wild-type levels of ExoN activity (Fig. 7 and 9). Based on nsp14 conservation (Fig. 1) and the viable phenotype of SARS-CoV and MHV ExoN-knockout mutants, MERS-CoV was expected to tolerate ExoN inactivation, in particular since the enzyme was proposed to improve the fidelity of CoV replication without being essential for RNA synthesis *per se* (10, 14, 21–23, 26, 29). This notion is further supported by the fact that the CoV RdRp (nsp12) exhibits *in vitro* activity in the absence of nsp14 (27). We therefore anticipated that an excess of deleterious mutations would first have to accumulate before becoming detrimental to MERS-CoV viability. Contrary to these expectations, an immediate and complete block of RNA synthesis was observed when MERS-CoV ExoN knockout mutants were launched by transfection of full-length RNA transcripts. It is noteworthy that similar observations were previously made for the corresponding ExoN knockout mutants of the alpha-CoVs HCoV-229E (19) and TGEV (34), the gamma-CoV avian infectious bronchitis virus (IBV) (E. Bickerton, S. Keep, and P. Britton, personal communication), and – according to a recent report that awaits peer review - also for SARS-CoV-2 (30), a close relative of SARS-CoV.

None of the ExoN mutations tested had a negative effect on the *in vitro* activity of the N7-MTase domain of nsp14, which is deemed essential for viral mRNA capping (Fig. 10). This is consistent with previous observations for SARS-CoV nsp14, in which the ExoN and N7-MTase activities were shown to be functionally distinct, though structurally interconnected by a hinge region that confers flexibility (21, 36). Given their unchanged N7-MTase activity, the non-viable phenotype of MERS-CoV ExoN active site mutants must be attributed to a negative effect on an additional and apparently critical function of the ExoN domain, which is directly involved in primary RNA synthesis rather than in (longer-term) fidelity control. At present we cannot explain, why SARS-CoV and MHV ExoN knockouts can apparently tolerate ExoN active site substitutions that are lethal to five other CoVs (now including MERS-CoV). Within the betacoronavirus group, in our experience, SARS-CoV and MHV display the most robust RNA synthesis and replication in cell culture when compared to MERS-CoV and SARS-CoV-2 (68–70). Possibly, the recovery of viable progeny depends on reaching a minimum level of RNA synthesis, which may somehow be achieved by only the most efficiently replicating CoVs. Admittedly, even bearing this possibility in mind, it remains difficult to reconcile the 1-to 2-log reduction of progeny titers observed for MHV and SARS-CoV ExoN-knockout mutants with the complete loss of infectious progeny observed for the ExoN-knockout mutants of the other CoVs.

In order to eliminate technicalities that might somehow prohibit the successful recovery of MERS-CoV ExoN knockout mutants and explain the phenotypic differences with other CoVs, we explored various details in the transfection protocol. This included the use of a DNA-launched system, similar to that used for TGEV (34), and the propagation of progeny virus in both innate immune-competent and - incompetent cells (Huh7 and Vero cells, respectively). However, this did not change the negative outcome of our transfection experiments, which were repeated more than 10 times for several mutants, always using wt and E191D MERS-CoV as positive controls that proved to be consistently viable. Our extensive set of MERS-CoV ExoN mutants tested, and the results obtained with various other CoVs (see above), strengthen our conclusion that – in addition to its proposed role as a proofreading enzyme – ExoN must have another function in CoV RNA synthesis (57).

As reported for MHV and SARS-CoV ExoN mutants (23, 71), possible (late) reversion was observed for a few of our MERS-CoV ExoN active site mutants, specifically mutants E191A, D273E, and in particular D90E, which had reverted at 6 d p.t. in three out of six experiments. This suggests that these mutants exhibit a low residual level of RNA synthesis that is the basis for these low-frequency events. In follow-up studies with the crippled H229C ZF1 mutant, a possible pseudo-revertant carrying a second-site mutation (Q19R) in nsp8 was identified in three independently obtained progeny samples, providing genetic support for an interaction between nsp8 and nsp14, which may be relevant in the context of the association of nsp14 with the tripartite RdRp complex consisting of nsp7, nsp8, and nsp12 (21, 27, 72–74). Future studies will address the properties of these nsp14 ExoN knockout mutants and their (pseudo) revertants in more detail.

In the only viable MERS-CoV ExoN active site mutant obtained, E191D, the catalytic motif was changed from DEEDh to the DEDDh that is characteristic of all members of the exonuclease family that ExoN belongs to (14, 15, 75). The phenotype of the E191D virus mutant was comparable to that of wt virus (Fig. 4). Biochemical assays revealed that E191D-ExoN enzyme is able to hydrolyze a dsRNA substrate with an activity level approaching that of the wt protein (Fig. 9). Additionally, the E191D mutant behaved similar as wt MERS-CoV upon treatment of infected cells with the mutagenic agent 5-FU (Fig. 4C-D) (28, 71), again suggesting its ExoN activity is not dramatically altered by this conservative substitution in the active site.

In this study, we developed an *in vitro* assay to evaluate MERS-CoV ExoN activity using a largely double-stranded RNA substrate (Fig. 6 and 7). As previously observed for SARS-CoV nsp14 (26), MERS-CoV ExoN activity was strongly enhanced by the presence of nsp10 (Fig. 8), in line with the formation of an nsp10:nsp14 heterodimer as observed in biochemical and structural studies (20, 26, 76). Slightly different patterns of degradation of the H4 substrate were observed when comparing the SARS-CoV and MERS-CoV ExoN enzymes *in vitro*. Likewise, the exchange of the nsp10 co-factor for the nsp10 subunit of the other virus, using the same substrate and the same nsp10:nsp14 ratio (1:4; Fig. 8), resulted in a slightly different pattern of substrate degradation, suggesting minor differences in the interaction of the nsp10:nsp14 complex with this particular RNA substrate. Previously, it was demonstrated that nsp10 is interchangeable between CoV subgenera in its role as co-factor for the nsp16 2′-O-methyltransferase, which was attributed to the high level of conservation of the nsp10-nsp16 interaction surface (77). As nsp14 and nsp16 share a common interaction surface on nsp10 (21, 26, 66), we explored whether a similar co-factor exchange was possible in the context of nsp14’s ExoN activity, which was indeed found to be the case (Fig. 8). Structurally, nsp14 interacts with nsp10 figuratively similar to a “hand (nsp14) over fist (nsp10)” conformation (21). In the formation of this complex, nsp10 induces conformational changes in the N-terminal region of ExoN that adjusts the distance between the catalytic residues in the back of the nsp14 palm and, consequently, impact ExoN activity (21). The exchange of the nsp10 co-factor between the two beta-CoVs might affect this conformation and, consequently, modulate the ExoN activity of the nsp14:nsp10 complex.

Alanine substitutions of active site residues severely reduced but did not completely abrogated the *in vitro* activity of MERS-CoV ExoN (Fig. 7-9), as previously shown for certain SARS-CoV nsp14 mutants (20, 36). Based on the two-ion-metal catalytic mechanism underlying the exonuclease activity of DEDDh family members (17, 56) and the SARS-CoV nsp14 structure, it was predicted that the various ExoN motifs contribute differently to the excision of nucleosides monophosphates (20, 21). Mutation of ExoN catalytic residues can alter ion binding (31) or disturb the fragile chemical equilibrium, as shown for conservative mutations (corresponding to E191D and D273E) in the Klenow fragment, a member of the DEDDh exonuclease family, which reduced ExoN activity by >96% (78). In general, all DEEDh mutations that yielded non-viable virus mutants exhibited similarly low levels of residual ExoN activity *in vitro* (Fig. 9), indicating that each of these residues is important for catalysis.

Our study suggests that, in addition to the active site residues, also other motifs in MERS-CoV ExoN are important for virus viability, specifically the two ZF motifs that were probed using two point mutations each (Fig. 2A). In previous ZF1 studies, a mutation equivalent to H229A created solubility issues during expression of recombinant SARS-CoV nsp14 (20) and resulted in a partially active ExoN in the case of white bream virus, a torovirus that also belongs to the nidovirus order (79). It was suggested that ZF1 contributes to the structural stability of ExoN, as it is close to the surface that interacts with nsp10 (20). Here, we demonstrate that the more conservative H229C replacement, which converts ZF1 from a non-classical CCCH type ZF motif into a classical CCCC type (63), was tolerated during recombinant protein expression and yielded an ExoN that is quite active *in vitro* (Fig. 7). This likely contributed to the fact that the H229C virus mutant retained a low level of viability (Fig. 2), although its overall crippled phenotype and the non-viable phenotype of mutant C201H clearly highlight the general importance of ZF1 for virus replication. In contrast, the corresponding TGEV mutant (ZF-C) was not strongly affected and could be stably maintained over several passages (34) The reverse genetics data suggest that ZF2, which is in close proximity to ExoN catalytic residues (20), is equally important, although technical complications with expression of the C261A and H264R nsp14 mutants prevented us to perform *in vitro* activity assays.

Like the ExoN domain of the arenavirus nucleoprotein (32, 33), the CoV ExoN was proposed to be involved in innate immune evasion (34, 35, 80), possibly by degrading viral ds RNA that in the case of CoVs is confined to characteristic double-membrane vesicles (81, 82). For TGEV, this suggestion was based on the reduced accumulation of dsRNA by the ZF-C mutant, which however remains to be characterized in more detail. In the absence of a TGEV ExoN activity assay, and in view of our data for the equivalent MERS-CoV ZF1 mutant, it seems premature to assume that the reduced levels of dsRNA in infected cells are caused by increased exonuclease activity of the ZF-C ExoN mutant (34).

In general, the properties of viable CoV ExoN mutants warrant further analysis. In future studies, the repertoire of residues probed by site-directed mutagenesis should be extended beyond active site and ZF motifs, which may help in particular to establish how directly reduced ExoN activity, primary viral RNA synthesis and enhanced innate responses are connected. Regardless of its possible interactions with host cell pathways, nsp14 clearly is a key subunit of the multi-enzyme complex that drives CoV genome replication, subgenomic RNA synthesis, and RNA recombination. Understanding the structure-function interplay between ExoN and other (viral and/or host) components will be key to elucidating its role in CoV RNA synthesis and evolution (83, 84). Taking into account the current SARS-CoV-2 pandemic, understanding the phenotypic differences between ExoN knockout mutants of different CoVs may contribute to the design of improved antiviral approaches, including those relying on ‘lethal mutagenesis’ or direct interference with viral RNA synthesis.

## Materials and methods

### Cell culture

Baby hamster kidney cells (BHK-21; ATCC CCL10), Vero and HuH7 cells were cultured as described previously (69, 85). Vero cells were kindly provided by the Erasmus Medical Center, Rotterdam, the Netherlands and HuH7 cells by Dr. Ralf Bartenschlager from Heidelberg University. For transfections, cells were maintained in Eagle’s minimal essential medium (EMEM; Lonza) with 8% fetal calf serum (FCS; Bodinco) supplemented with 100 IU/ml of penicillin and 100 μg/ml of streptomycin (Sigma), and 2 mM L-Glutamine (PAA), and incubated at 37 °C with 5% CO2. Infection of Vero, Vero E6 and HuH7 cells was carried out in EMEM containing 2 % FCS.

### Reverse genetics

Mutations in the MERS-CoV nsp14-coding region were engineered in a bacterial artificial chromosome (BAC) vector (51, 52) containing the full-length cDNA copy of MERS-CoV strain EMC/2012 (44, 53), by two-step e*n passant* recombineering in *E. coli* (86). When designing the primers, a translationally silent marker mutation was introduced near the site of mutagenesis in order to differentiate between the occurrence of reversion and (possible) contamination with parental virus. For each mutation, two mutant BACs were isolated independently, the nsp14-coding region was verified by sequencing, and both BACs were used for *in vitro* run-off transcription and virus launching.

Approximately 5 μg of BAC DNA was linearized with *Not*I and full-length RNA was obtained by *in vitro* transcription with T7 RNA polymerase followed by lithium chloride precipitation according to the manufacturer’s protocol (mMessage-mMachine T7 Kit; Ambion). 5 μg of RNA was electroporated into 5×10^6^ BHK-21 cells using the Amaxa nucleofector 2b (program A-031) and Nucleofection T solution kit (Lonza). Transfected BHK-21 cells were mixed with HuH7 or Vero cells in a 1:1 ratio and plated for harvesting supernatants, intracellular RNA isolation and analysis by immunofluorescence microscopy. Immunolabelling was performed as described before (69), using antibodies recognizing double-stranded RNA (dsRNA; (87)), M protein (52) or a cross-reacting antibody raised against SARS-CoV-nsp3 (88). Cells were incubated at 37⁰C up to a maximum of 6 days post transfection (d.p.t.). Supernatants were collected when full cytopathic effect was observed or at the end of the experiment. Virus titers were determined by plaque assay in HuH7 and Vero cells (89). In order to confirm the presence of engineered mutations in viral progeny, HuH7 and Vero cells were infected with supernatants harvested from transfected cells and intracellular RNA was isolated at 18 h post infection as described above. cDNA was synthesized by reverse transcription using RevertedAid H minus reverse transcriptase (ThermoFischer Scientific) and random hexamer primers (Promega), in combination with a primer targeting the 3’ end of the MERS-CoV genome. The full-length genome or the nsp14-coding region were amplified by PCR using MyTaq DNA polymerase (Bioline) and after purification the PCR product was sequenced by Sanger sequencing. Genome sequencing by NGS was performed as described before (70). All work with live recombinant MERS-CoV was done in a biosafety level 3 laboratory at Leiden University Medical Center.

### Analysis of viral RNA synthesis

Isolation of intracellular RNA was performed by lysing infected cell monolayers with TriPure isolation reagent (Roche Applied Science) according to the manufacturer’s instructions. After purification, intracellular RNA samples were loaded onto a 1.5% agarose gel containing 2.2 M formaldehyde, which was run overnight at low voltage overnight in MOPS buffer (10 mM MOPS (sodium salt) (pH 7), 5 mM sodium acetate, 1 mM EDTA). Dried agarose gels were used for direct detection of viral mRNAs by hybridization with a ^32^P-labeled oligonucleotide probe (5’-GCAAATCATCTAATTAGCCTAATC-3’) that is complementary to the 3’-terminal sequence of MERS-CoV genome and all subgenomic mRNAs. After hybridization, RNA bands were visualised (using exposure times of up to 28 days) and quantified by phosphorimaging using a Typhoon-9410 variable mode scanner (GE Healthcare) and ImageQuant TL software (GE Healthcare).

PCR primers and Taqman probes targeting ORF1a (junction of nsp2-nsp3 coding region), the nucleocapsid (N) protein gene, or the leader-body TRS junction of subgenomic mRNA3 were designed and analyzed for multiplex quality using Beacon Designer™ Software (Premier Biosoft). Reverse transcription (RT) was performed using RevertedAid H minus reverse transcriptase (ThermoFischer Scientific) and a mix of specific reverse primers targeting ORF1a, ORF8, or subgenomic RNA 3 (primer sequences used available upon request). The mRNA derived from the cellular β-actin gene was used as a reference housekeeping gene. Tagged primers were used to differentiate between positive- and negative-stranded viral RNA. Samples were assayed by Taqman multiplex real-time PCR using TaqMan Universal Master Mix II and a CFX384 Touch™ Real Time PCR detection system (BioRad). A standard curve was obtained using an *in vitro* transcript derived from a synthetic plasmid that contained all PCR targets. cDNA was obtained as described above. Each RNA sample was analyzed in triplicate.

### Plaque reduction assay

HuH7 cells seeded in 6-well clusters were infected with recombinant MERS-CoV at low MOI (30 PFU/well) for 1 h at 37°C. Subsequently, the inoculum was replaced with 2 ml of a 1.2% suspension of Avicel (RC-581; FMC Biopolymer (90) in DMEM (containing 2% FCS and antibiotics) and serial dilutions of 5-FU (F6627, Sigma-Aldrich) or Ribavirin (R9644, Sigma-Aldrich) ranging from 0 to 400 μM. Cells were incubated at 37°C for 72 h, fixed with 7.4% formaldehyde, and plaques were visualized using crystal violet staining.

To compare the effect of 5-FU treatment on the progeny titers of wt and nsp14-E191D rMERS-CoV, confluent monolayers of HuH7 were incubated for 30 min at 37⁰C with solvent or a range of 5-FU concentrations. The drug was then removed and cells were infected at an MOI of 0.1 during 1 h at 37⁰C. After removal of the inoculum, EMEM containing 2% FCS and solvent or a matching concentration of 5-FU was added to the wells. Supernatants were collected after 30h and rMERS-CoV progeny titers were determined by plaque assay. All drug-treated samples were normalized to the untreated vehicle control, and values were expressed as fold change compared to untreated virus titers.

### Expression and purification of recombinant CoV nsps

SARS-CoV nsp10 and nsp14 were produced as described before (25) and used as a positive control in all biochemical assays. A MERS-CoV nsp10 expression construct was kindly provided by Dr. Deyin Guo (77) (91) and purified according to their protocol. All MERS-CoV nsp14 constructs were cloned into expression vector pDEST14 with an N-terminal His6-tag using the Gateway system (25). MERS-CoV nsp14 mutant expression plasmids were generated by Quikchange site-directed mutagenesis using Accuzyme DNA polymerase (Bioline) following the manufacturer’s instructions. pDEST14 plasmids expressing MERS-CoV nsp14 were transformed into competent *E. coli* strain Rosetta (DE3) pLyS (Novagen) and cultured in Luria-Bertani (LB) broth supplemented with 100 μg/ ml of ampicillin and 30 μg/ ml of chloramphenicol. Protein expression was induced at an optical density (OD600nm) of 0.8 by adding 50 μM of isopropyl-β-D-a-thiogalactopyranoside (IPTG; Bioline). After 24 h at 13⁰C, induced cells were harvested and lysed in a buffer containing 50 mM Tris-HCl, pH 7.5, 150 mM NaCl, 5 mM β-mercaptoethanol, 5% glycerol, 1 mM PMSF, and 20 mM imidazole (92). Next, the lysate was centrifuged at 12,000xg for 30 min, and the soluble fraction was column-purified by immobilized metal ion affinity chromatography using Nickel sepharose high performance beads (17526802, GE Healthcare). The eluate was fractionated by gel filtration on a Superdex-200 Increase 10/300GL column (GE Healthcare) in buffer containing 30 mM HEPES, pH 7,5, 300 mM NaCl, and 5% glycerol. In the end, proteins were concentrated using ultrafiltration devices with a molecular mass cut-off of 30 kDa (Millipore), and protein concentrations were measured using spectrophotometry. All purified proteins were analyzed by SDS-PAGE followed by Coomassie blue staining as well as by Western blot using a mouse monoclonal antibody against the 6xHis-Tag (Novagen). Protein aliquots were stored at −80⁰C in 50% glycerol (v/v) and used for enzymatic assays.

### Exonuclease activity assay

Synthetic RNA H4 (26) was radiolabeled at its 5’ end using T4 polynucleotide kinase (Epicentre) and [γ-^32^P]ATP (Perkin Elmer). Unless stated otherwise in figures or legends, reactions contained 200 nM of recombinant nsp14, 800 nM of nsp10, and 750 nM of radiolabeled substrate in 40 mM Tris-HCl pH 7.5 containing 5 mM of MgCl2 and 1 mM of DTT. After incubation at 37⁰C for up to 90min, reactions were stopped by addition of an equal volume of loading buffer containing 96% formamide and 10mM EDTA. Samples were then loaded on 7M urea-containing 20% (wt/vol) polyacrylamide gels (acrylamide/bisacrylamide ratio 19:1) buffered with 0.5x Tris-borate-EDTA and run at high voltage (1600 V). Results were visualized by phosphorimaging as described above.

### N7-methyltransferase activity assay

Methyltransferase assays were performed in 40 mM Tris-HCl, pH 8.0, 5 mM DTT, 2 μM of ^7Me^GpppA or GpppA RNA cap analogue (New England Biolabs), 10 μM adenosyl-methionine (AdoMet, Thermofisher), 0.03 μCi/μl [^3^H]AdoMet (PerkinElmer) (25). In each reaction, MERS-CoV or SARS-CoV nsp14 was added to a final concentration of 500 or 250 nM, respectively. Reactions were incubated at 30⁰C for up to 120min, and were stopped by the addition of 10-fold volume of 100 μM ice-cold Adenosyl-Homocysteine (AdoHcy; ThermoFischer). Then, samples were spotted on a DEAE filtermat (PerkinElmer) pre-wet with Tris-HCl pH 8.0 buffer. Filtermats were washed twice with 10 mM ammonium formate (Sigma-Aldrich), pH 8.0, twice with MiliQ water, and once with absolute ethanol (Sigma-Aldrich). After air drying for 10 min, filtermats were cut and relevant pieces transferred to individual tubes. Betaplate scintillation fluid (PerkinElmer) was added and the amount or ^3^H-label bound was measured in counts per minute (cpm) using a Wallac scintillation counter. For relative quantification, incorporation measurements for mutant proteins were normalized to values obtained with the wt control nsp14. Samples were measured in duplicate in each experiment.

## Acknowledgments

N.S.O. was supported by the Marie Skłodowska-Curie ETN European Training Network ‘ANTIVIRALS’ (EU Grant Agreement No. 642434). C.C.P and E.J.S. were supported in part by TOP-GO grant 700.10.352 from the Netherlands Organization for Scientific Research. We thank Yvonne van der Meer for technical support and Linda Boomaars for providing SARS-and MERS-CoV nsp10 and nsp14 expression constructs. We are grateful to Etienne Decroly, Francois Ferron and Bruno Canard (University of Aix-Marseille, France) for helpful discussions. We kindly acknowledge the sharing of unpublished information on IBV ExoN knockout mutants by Dr. Erica Bickerton and colleagues (Pirbright Institute, U.K.).

